# Site, Fertilization and Season Structure the Soil Microbiome and its Interactions with *Bdellovibrio* and Like Organisms Predators

**DOI:** 10.64898/2026.06.29.735237

**Authors:** Alka Kumari, Rolf Lood, Ofra Matan, Eddie Cytryn, Yael Laor, Gil Eshel, Edouard Jurkevitch

## Abstract

The contribution of predation between bacteria to microbial community dynamics in agricultural fields has hardly been investigated. Here. dynamics of general prokaryotes (GEP) and of the predators Bdellovibrionales (Bd) and Bacteriovoracales (Bac) (*Bdellovibrio*-and-Like Organisms, BALOs) were studied in two agricultural fields differing in organic and mineral input regimes, for one year. Season, but not fertilization, affected absolute sizes of GEP and of BALO communities. 16S rRNA gene community analysis identified numerous novel Bd and Bac lineages, with none of the dominant BALOs related to characterized isolates. A few dominant BALO amplicon sequence variants (ASVs) persisted year-round, whereas others showed seasonal- or treatment specific responses. GEP, Bd, and Bac ASV a-diversity was mostly influenced by season, with some changes due to fertilization in Bd, and Bac communities. Seasonal changes, site, and fertilization regimes influenced β-diversity of GEP, Bd and Bac communities and determined the structure of BALO-gram-negative bacteria interaction networks, signaling that niche segregation acts at the microbiome-BALO interface. Accordingly, we suggest that shifts in GEP community structure triggered by environmental changes and agricultural practices cascade to BALO predators, in turn affecting BALO-microbiome interactions. These dynamics may be harnessed to manipulate the soil microbiome to benefit sustainable environmental and agricultural outcomes.

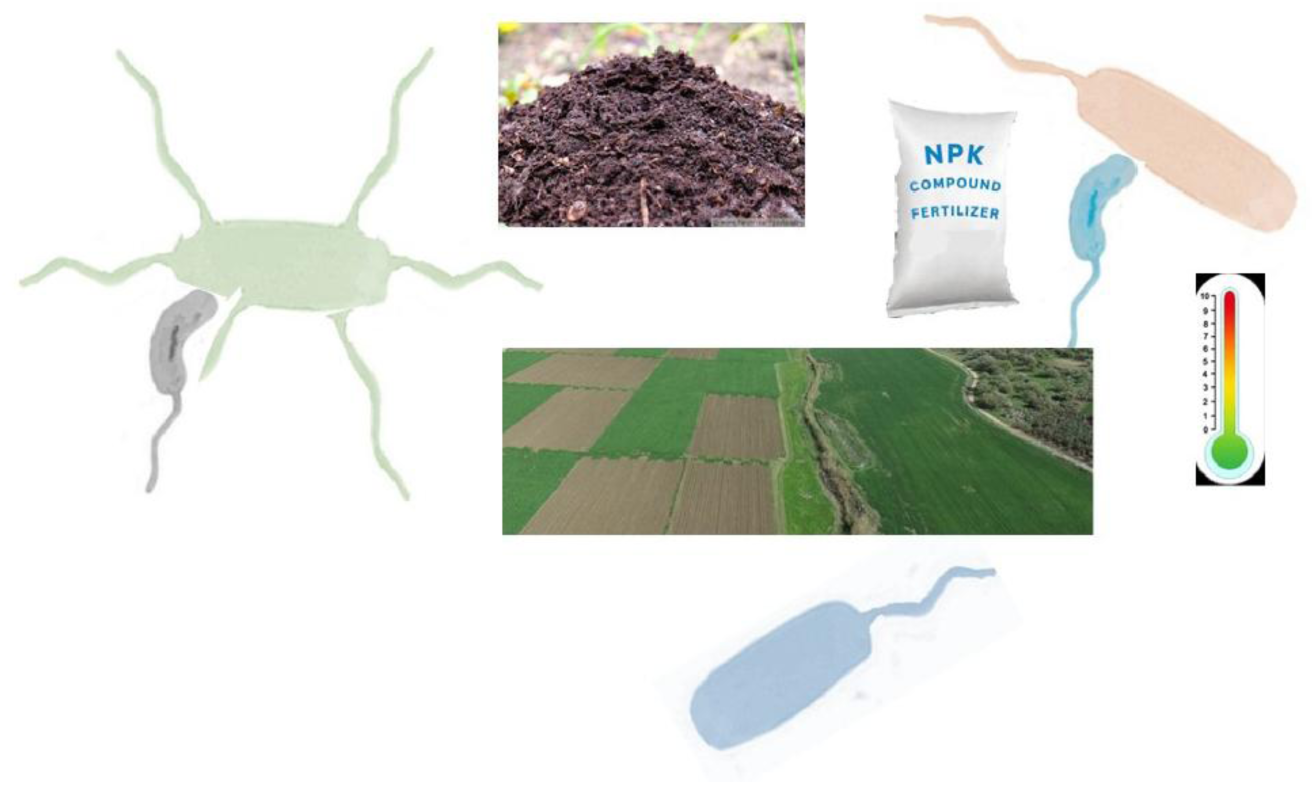

The soil microbiome and the microbial predators *Bdellovibrio* and Like Organisms (BALOs) are influenced by environmental parameters (season) and agricultural practice (fertilization) in their population dynamics, diversity and interactions. BALO diversity is high, and novel clades are found with BALOs differing in abundance and persistence. Microbiome-BALO interactions suggest environment and agricultural practice-dependence of niche segregation in BALOs. BALOs – and other micro-predators- may thus be integrated in the development of soil microbiome manipulations for environmental and agricultural applications.

## Introduction

Intensive agriculture (thereafter “conventional agriculture, CA”), while essential for food production and security, generates large scale environmental damage ^1,2^. Chemical fertilizers provide immediately available nutrients able to sustain high concentrations in the soil solution and boost plant growth ^3,4^. They are therefore intensively used, making them one major cause of pollution of soils and of water sources, and their production an important source of greenhouse gases ^5–7^. Moreover, their exclusive use as sources of plant nutrients depletes soils of organic matter, resulting in tremendous losses in soil organic carbon (SOC) which in turn, increases soil erosion and may exacerbate climate change ^8^. The positive results of CA on crop production are thus jeopardized by negative effects on soil health including physicochemical properties and long-term fertility, environmental pollution and loss of biodiversity ^8,9^. Organic farming (OF) are low-(external) input agro-ecosystems that utilize the availability of nutrients naturally found in plant and animal-derived material to sustain productivity and ecosystem functionality as well as to provide control of biological pathogens^10^. The large amounts of added organic material in OF contributes to SOC sequestration, reduces erosion by stabilizing soil aggregates and improving aggregate structure, and promotes water conservation ^11^. In particular, OF enhances microbial activity as microbial populations slowly decompose organic amendments and release nutrients that contribute to soil stabilization, long-term fertility and health, and increase above and below ground biological diversities that in turn further enhance these beneficial soil properties ^3,9,12–16^. OF reduces the ecological footprint of agriculture, providing a more sustainable alternative to CA^17^. It has been shown that these changes in the soil habitat bring about shifts in the soil microbiome, although different studies can differ in the type and strength of the effects; For example, in measures like abundance of general bacterial or specific phyla, diversity and heterogeneity, enzymatic activities, and specific functional traits like phosphate solubilization, nitrogen fixation or cellulose degradation ^9,18–22^. Although numerous studies have shown that soil management and environmental heterogeneity influence microbial composition and function, less have explored how seasonal changes and fertilization regimes modulate interaction dynamics within microbial communities.

Microbiome functionality studied through trophic networks show strong effects of predatory interactions on network structure ^23,24^. A meta-analysis on the interactions of protists with bacterial communities showed that predatory protists dominate the overall microbial community in soil, and that their richness and relative abundance positively correlated with bacterial richness in forest and grassland soils, but not in agricultural soils ^25^. These finding suggests that higher microbial community trophic levels may be affected by agricultural land management, altering trophic complexity and ecosystem functions ^25^. While chemical fertilization decouples this trophic network, OF supports protistan predation which may subsequently contribute to improved soil health and control the spread of plant pathogens ^26,27^.

Myxobacteria (order *Myxococcales*) are common, widely abundant facultative, soil-dwelling bacterial predators with broad prey ranges ^28–30^. *Bdellovibrio* and Like Organisms (BALOs; mainly orders Bdellovibrionales and Bacteriovoracales) are obligate predators widely distributed in the environment ^31^. Both groups have a preponderant role in regulating nutrient flow in soil ^32,33^. Specifically BALOs were shown to influence bacterial community composition, nutrient turnover, and help suppress pathogens ^33–36^, and respond to fluctuations in the abundance of specific populations in rice paddies ^37^. In the rhizosphere as well as in wastewater, BALO populations are diverse and vary in prey range ^38,39^. Despite their ecological significance, studies on BALO dynamics in soil, and particularly in agricultural soil remain limited.

In this study, we aimed to address this knowledge gap. We hypothesized that soil and crop management practices impact the abundance, composition, and trophic dynamics of the soil microbiome and the BALOs predators associated to it. In that order, we obtained 16S rDNA high-throughput amplicon sequence variant (ASVs) and PCR quantitative data of bacteria and predator communities in two agricultural fields for one year and measured the effects of organic and inorganic fertilization and of environmental parameters on their abundances, community structures, and interactions.

## Materials and methods

### Experimental sites and treatments

Two agricultural fields at the Newe Ya’ar, Research Center of the Israel Agricultural Research Organization-Volcani Institute were examined (Figure S1, Table S1). At the Model Farm (MF) field, two amendments were applied in October 2020: Pre-stabilized cow manure brought from a local dairy farm (Bet She’arim), at a rate of 22 m³ ha^-1^, and manure-based compost brought from an industrial facility (’Ha’amakim’), at 35 m³ ha^-1^, the latter produced from livestock manure and slaughterhouse blood residues. Corn was sown in early April 2022 and soil sampling campaigns launched thereafter at four time points to capture seasonal variation: April (spring) 2022, immediately after sowing; July (summer) 2022, during crop growth when the fields were irrigated with treated wastewater; November (Autumn) 2022, following harvest and ploughing, with residual plant roots left in the soil; and February (winter) 2023, during the fallow winter period (Figure S2). The Organic Plots (OP) field is managed under organic farming since 2009 ^40^ and is part of a typical organic rotation system at Newe Ya’ar station ^41^. At the end of summer 2021, the field was treated with one of three fertilization regimes: mineral fertilizer (NPK at 200, 50, and 100 Kg ha^-1^), low-dose compost (10 m³ ha^-1^), or high-dose compost (60 m³ ha^-1^). The field was subsequently planted with broccoli in September 2021. The same fertilization treatments were reapplied in August 2022 before the following cultivation cycle. Soil samples were collected at two time points: April 2022, following the 2021 amendment application and prior to the August 2022 reapplication, and November 2022, after the second amendment application (Figure S2).

At both fields, soil was sampled from the top 10 cm layer using an auger, with sampling points positioned approximately 25 cm apart, or 25 cm from the plant stem when vegetation was present. The same GPS-referenced locations were used across all sampling events to ensure consistency. Each treatment was sampled at four randomly selected locations that were subsequently used as fixed sampling points across all time points (four biological replicates per treatment). This produced 32 samples from the MF site and 24 from the OP site. Prior to analysis, visible debris and stones were manually removed from all samples.

### Sample processing and DNA extraction

Soil DNA was extracted from the collected samples using the DNeasy PowerSoil Pro Kits following the manufacturer’s protocol (Qiagen, Germany). For each sample, 250 mg of soil was weighed and placed in an Eppendorf tube for extraction. DNA yield and purity were measured using a Nanodrop spectrophotometer (Thermo Fisher, USA). The extracted DNA samples were stored at -20°C until further analysis.

### Soil physicochemical properties

Soil pH was measured with a calibrated pH meter by saturating 10 g of soil with 10 ml of autoclaved double distilled water (DDW) with stirring to ensure homogeneity. Moisture content was determined by weighing samples before and after overnight drying at 95°C. Dried samples were then finely ground with a sterile mortar-mortar pestle for homogeneity. Approximately 10 mg of homogenized soil was used for total carbon (TC) and total nitrogen (TN) analysis by combustion analysis. Relative humidity and air temperature were measured at the site through visual crossing weather query builder (https://www.visualcrossing.com/) based on the GPS coordinates (Table S1).

### Bacterial quantification by real-time QPCR

Standard curves for Bdellovibrionaceae (qBd) and Bacteriovoracaceae (qBac) quantification were prepared using ten-fold serially diluted plasmids containing 1467 bp fragments of the 16S rRNA genes of *Bdellovibrio bacteriovorus* HD100 and of *Bacteriovorax stolpii* UKi2, respectively, cloned into pGEM-T Easy vectors (Promega, USA) and transformed in *E. coli* TG1 ^42^. *E. coli* ML21 was used for constructing a GEP standard curve. Primers for GEP, qBd, and qBac, and amplification protocols are listed in Table S2. Each 10 μl reaction consisted of 5 μl of SYBR® Green PCR Master Mix (Applied Biosystems, California, USA), 0.1 μl of each primer (10 μM), 1 μl of template DNA and 3.8 μl of PCR grade DDW.

### High-throughput 16S rRNA gene sequencing

16S rRNA gene amplicon sequencing was performed using general primers targeting the gene’s V3 region for General Prokaryotes (GEP), and primers specifically targeting the Bdellovibrionales (Bd) and Bacteriovoracales (Bac) 16S rDNA genes, following a two-stage protocol ^42,43^ (Table S2). All samples were combined and sequenced using a MiSeq platform (Illumina) at the Rush Genomics and Microbiome Core Facility, RUSH University, Chicago, USA. A ZymoBIOMICS™ Microbial Community DNA Standard (Zymo Research, Irvine, CA, USA) was used as a positive control, with sterile DNA-free water as the negative control.

### Sequence reads processing and analysis

Sequencing reads where processed according to the standard workflow of qiime2-2023.5 ^44,45^. Adapters and primers were removed from the paired-end sequence reads, further subjected to quality trimming using Cutadapt. Overlapping reads were merged and processed using DADA2 denoising. Non-chimeric Amplicon Sequence Variant (ASVs) were assigned a taxonomy based on reference sequences in SILVA release 138 ^46,47^. The resulting initial ASVs table was preprocessed for downstream analysis. Unclassified sequences at Kingdom level and non-bacterial reads (mitochondria, chloroplasts, and Eukarya) were removed from the dataset. Those ASVs constituted only by the mock samples and negative controls were removed. Finally, ASVs with fewer than 20 reads across the entire dataset were removed to obtain a fully processed ASV table.

### Microbial diversity and statistical analyses

α-diversity indices (Observed, Shannon and Simpson index) were calculated from ASVs count on a per-sample basis, using the Vegan R package. Comparison of α-diversity among different sample groups (season, fertilization type and sampling sites) was performed with the Wilcoxon test on the JMP statistical package (Carey, NC). Differences between microbiome structures were assessed using Bray-Curtis dissimilarity calculated on relativized datasets (according to sum of abundance in each sample). PCoA plots and Redundancy Analysis (RDA) were generated using the Vegan R package, with the first two principal coordinate axes explaining the greatest proportion of variance in the data. Comparisons of microbiome structure between groups were performed with PERMANOVA. Group-specific core microbiomes were identified as ASVs from GEP, Bd and Bac with >1% mean RA and present in ≥ 50% of samples within each treatment group. Similarity between core microbiomes was evaluated using Jaccard similarity indices of shared core ASVs.

### Phylogenetic analysis of BALOs

The top 100 most abundant Bd and Bac ASVs, based on relative abundance across soil samples were selected for multiple sequence alignment with MUSCLE (Multiple Sequence Comparison by Log-Expectation) ^48^. Gaps and poorly aligned positions were removed to create a set of unambiguously aligned sequences ^49^. The resulting aligned sequences of Bd (390 bp) and Bac (300 bp), including reference strains, were used to build an unrooted maximum likelihood (ML) tree using MEGA11 ^49^. A bootstrap consensus tree for each family was inferred from 100 replicates. Evolutionary divergence among Bd and Bac ASVs and their closest reference strains was estimated in MEGA11 ^51^ by calculating the number of base differences per site. The phylogenetic trees were annotated and visualized using in iTOL v5 ^50^.

### Potential interactions and co-occurrence network analysis

Since BALOs prey on Gram-negative bacteria, ASVs belonging to phyla Bacillota and Actinomycetota, and to domain Archaea were filtered out from the GEP dataset. This formed a “Gram-negative total bacteria” (GNtob) set used for the computation of individual co-occurrences and of networks with Bd and Bacs ASVs. GNtobs, Bd, and Bac ASVs from the fertilization treatments at the MF and OP sites were analyzed separately for each treatment. ASVs with less than 10 reads were filtered out to reduce complexity. Kendall rank correlations between GNtobs and Bd or Bac ASVs were calculated in each sample group. Negative correlations (τ < -0.3) were retained to represent interactions. Network edges were subsequently selected based on significant negative correlations (p < 0.05). Networks were visualized using the Fruchterman-Reingold force-directed layout implemented in Gephi (V0.10). Network topology was characterized by node degree (average number of GNtob ASVs connected to Bd or Bac ASVs), modularity index (degree of network subdivision into discrete ecological clusters), average node connectivity (interaction density across the network), and average path length (mean shortest distance between connected ASV pairs within the network) ^51^. Edge color represents the target predator group, while edge thickness corresponds to correlation strength (τ).

## Results

### GEP and BALO population dynamics in MF and OP fields

GEP abundance in MF field samples, as determined by QPCR (log copies g⁻¹) ranged from 8.2 ± 0.1 to 10.6 ± 0.05 and from 10.8 ± 0.03 to 9.3 ± 0.4 in compost and in manure-treated soils, respectively, decreasing between July and November (Figure 1A). qBd and qBac populations were both highest and of similar sizes in April under compost and manure treatments (Figure 1B, C). They decreased by two and one orders of magnitude in July, to increase in November and February, respectively. At the OP site under chemical, compost and high compost fertilization, GEP population sizes increased between April to November from 10.8 ± 0.1 to 11.8 ± 0.02. The qBd and qBac populations were of comparable sizes and similar to those at MF, in July, and in November (Figure 1D-F). Relative abundances (RA, % of GEP) of BALOs varied, being within the 0.1 to >1% range in almost all the MF site samples. In OP site samples, as a result of average higher GEP population sizes, BALO RAs remained <0.1%, (Figure S3).

**Figure 1.**
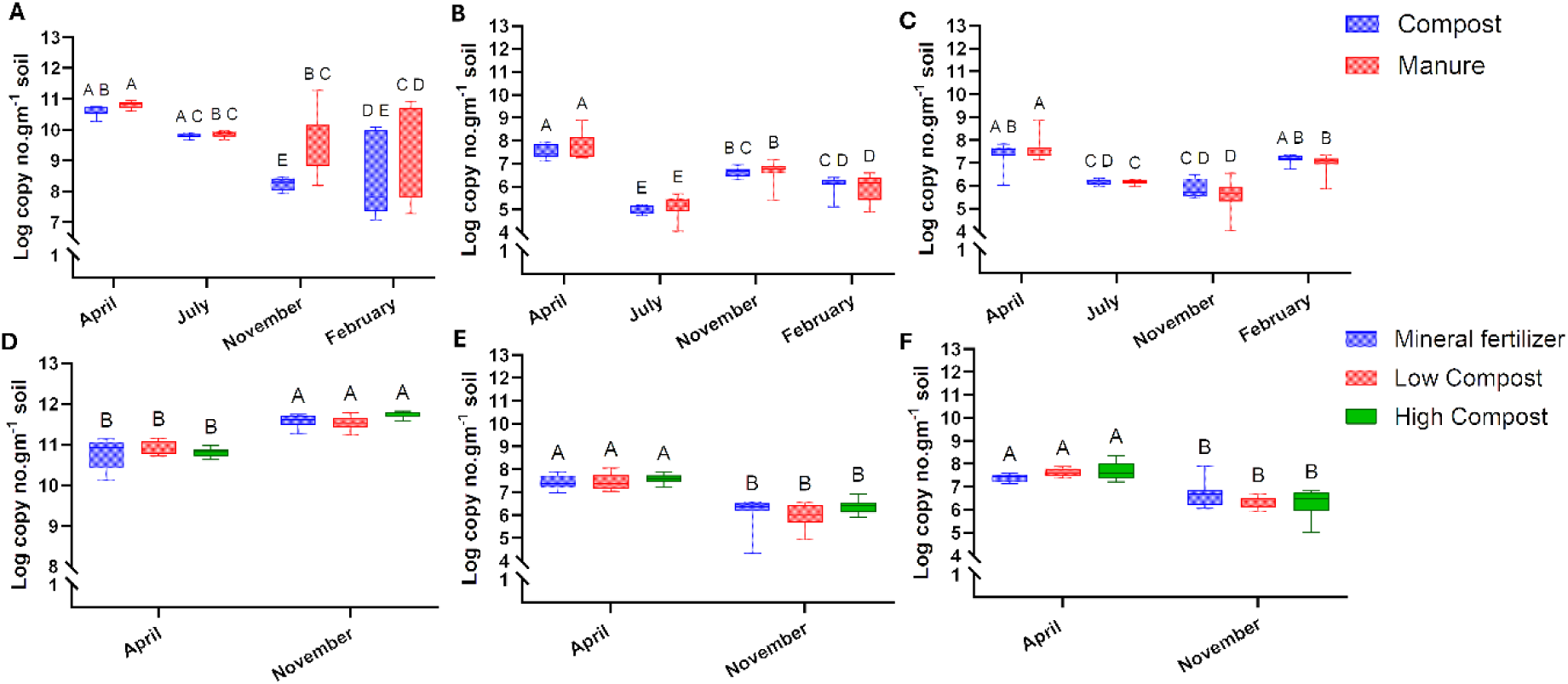
Quantitative PCR analysis of General Prokaryotes (A, D), Bdellovibrionaceae (B, E), and Bacteriovoracaceae (C, F) populations at the MF (A-C) and OP (D-F) sites. Different letters above the boxplots indicate statistically significant differences according to Tukey’s HSD post hoc test (p < 0.05).

### Summary of amplicon sequencing output

In total, 2,063,501, 3,539,836, and 4,186,675 raw reads for GEP, Bd and Bac, respectively were obtained, averaging 33,282 ± 11,086, 63,103.2± 13,288 and 74,727±52,690, respectively. Following processing, 24,361 ± 9,565 (55%), 26,998 ± 14,102.64 (41%), and 36,660 ± 41,526 (37%) reads were obtained for GEP, Bd and Bac, respectively, reaching saturation (Figure S4). After singleton and low-quality reads (<20) removal, this yielded 2402, 643 and 1314 non-chimeric ASVs for GEP, Bd and Bac respectively. Negative controls yielded on average 318 ± 167, 1009.5 ± 1498.9, and 385±299 reads for control DNA, PCR and sequencing steps of GEP, Bd and Bac, respectively.

### Site, fertilization, and seasonal effects on diversity

In MF field samples, GEP α-diversity remained stable in both (compost and manure) treatments in April and July, but decreased in November, and further declined in under compost amendment, whereas manure-treated soils maintained comparatively higher diversity (Table S3). GEP diversity at the OP site was comparable to that at MF at the same time points (April, November). At MF, diversity was highest in April (Bac) and April and July (Bd), followed by fluctuations in the following seasons, with differences between treatments (Tables S4, S5). In OP field samples, Bd richness largely increased in November (Table S4); whereas Bac diversity was very large in April and November in two of the three fertilization treatments (Table S5). PCoA ordination showed MF and OP samples formed visually distinct clusters (Figure 2A-C). GEP β-diversity was significantly structured by season (R² = 0.18, F = 4.47, p = 0.001, PERMANOVA), fertilization (R² = 0.13, F = 2.43, p = 0.001, PERMANOVA) and site (R² = 0.09, F = 6.69, p = 0.001, PERMANOVA). The Bd community was also structured by the same factors (Fertilization, R² = 0.21, F = 4.34, p = 0.001, PERMANOVA; Season, R² = 0.16, F = 4.58, p = 0.001, PERMANOVA, and; Site, R² = 0.14, F = 11.46, p = 0.001, PERMANOVA). For the Bac community, fertilization (R² = 0.12, F = 1.79, p = 0.001, PERMANOVA) contributed to β-diversity more than season (R² = 0.08, F = 1.65, p = 0.004, PERMANOVA) or site (R² = 0.04, F = 2.26, p = 0.001, PERMANOVA) (Figure 2C). RDA revealed that temperature, moisture, and humidity significantly influenced GEP composition, explaining 17.9% of total variation (Table S6, Figure S5A). The same parameters, and humidity, moisture and pH, explained 23.8 and 11.1% of the variance across samples of the Bd and Bac communities, respectively (Table S7, S8; Figure S5B, C).

**Figure 2.**
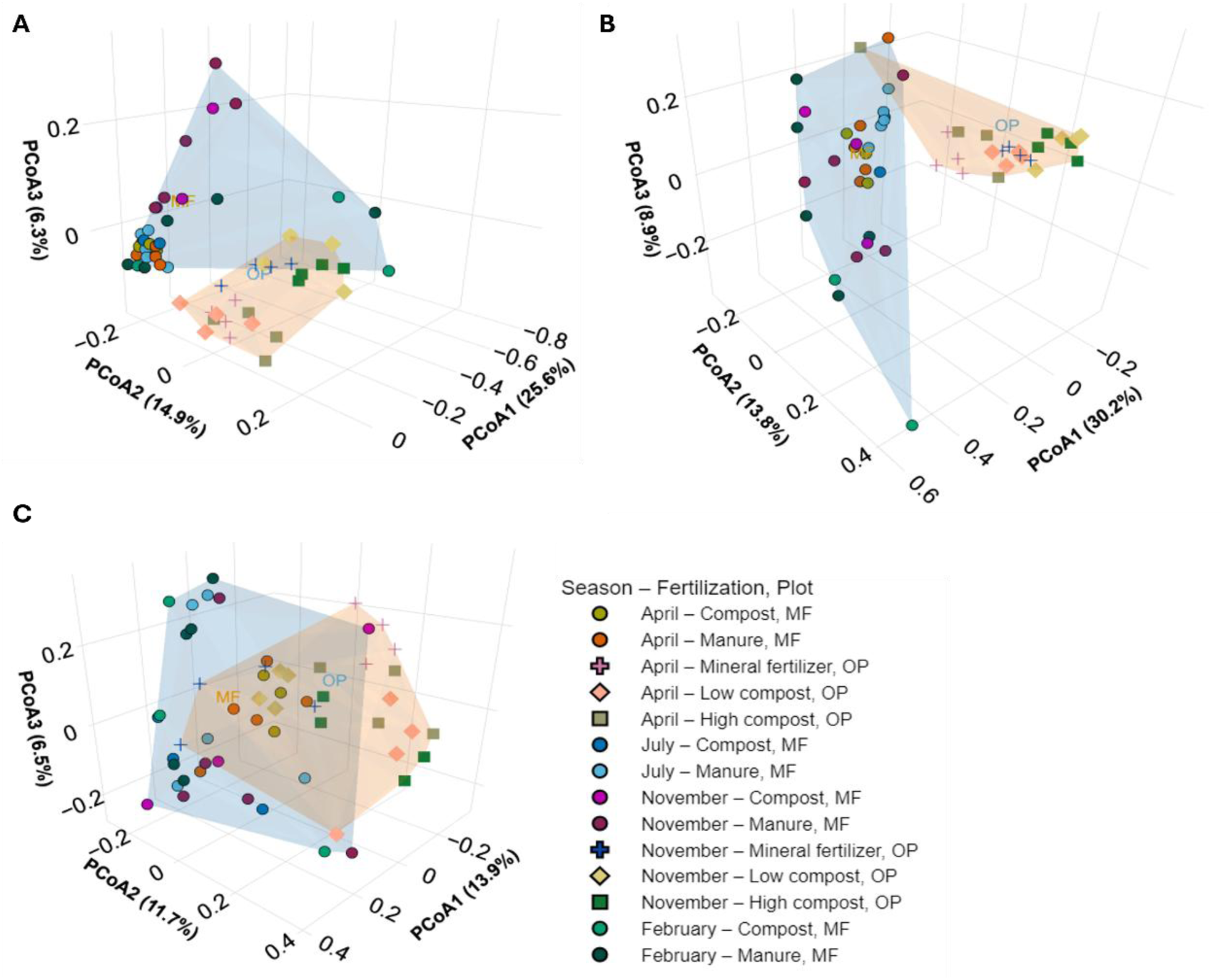
Principal Coordinates Analysis (PCoA) of soil microbiome based on Bray-Curtis dissimilarity matrix of General Prokaryotes (GEP) (A), Bdellovibrionales (B), and Bacteriovoracales (C) community distribution across sampling sites (MF, OP) and seasons. Convex hulls indicate site-specific clustering, with blue and beige hulls representing the MF the OP sites, respectively.

### Community composition and cores

Community composition of the GEP at the family level was largely consistent across treatments and seasons at both the MF and OP sites, with moderate shifts in relative abundance (RA). Core assemblage comprised the families Rubrobacteriaceae (7.6-11.0%), Solirubrobacterales (4.0-8.9%) and Planococcaceae (2.1-6.0%), followed by Beijerinckiaceae (2.1-5.5%), Cellulomonadaceae (1.9-3.5%), Gemmatimonadaceae (1.8-3.3%), Bacillaceae (1.3-3.8%) and Vicinamibacteraceae (3.5-6.2%). A substantial fraction of the taxa (18.6-34.5%), with mean relative abundance <1% (rare taxa) were pooled as “Others (Figure S6). At the genus level, the prokaryotic community was dominated by a limited number of genera alongside a large proportion of low-abundance and unclassified groups. Among classified genera, *Rubrobacter* (2.3-4.9%), *Microvirga* (1.4-4.4%), *Arthrobacter* (1-3.0%), *Gaiella* (0.9-2.6%), and *Bacillus* (0.8-2.5%) were consistently detected across samples (Figure S7).

The GEP core community showed high temporal persistence at both sites except in February in the MF field (Figure S8: Table S9); Nearly half of core GEP ASVs were shared between sites (253 ASVs - 49%) but the proportions of shared core predators ASVs between sites were lower (Figure S9; Table S10). The distributions of core Bd and Bac ASVs were biased, with their majority, and almost their totality, respectively, detected in OP samples (Figure S9; Table S10).

### BALO phylogeny

A maximum likelihood analysis was performed using the 100 most abundant Bd and Bac ASVs, of an observed total of 643 and 1314, respectively. In the Bd tree (Figure 3A), three major clusters were identified. Four of the five most abundant Bd ASVs were phylogenetically distinct from the known reference strains *Bdellovibrio bacteriovorus* HD100 and *B. reynosensis* LBG001. The Bac clustered as two major clades (Figure 3B). One cluster grouped with type strains *Bacteriovorax stolpii* DSM 12718 and *Halobacteriovorax marinus* JSS, while the second cluster diverged from known references. The top five most abundant Bac ASVs were distributed between both clades.

**Figure 3.**
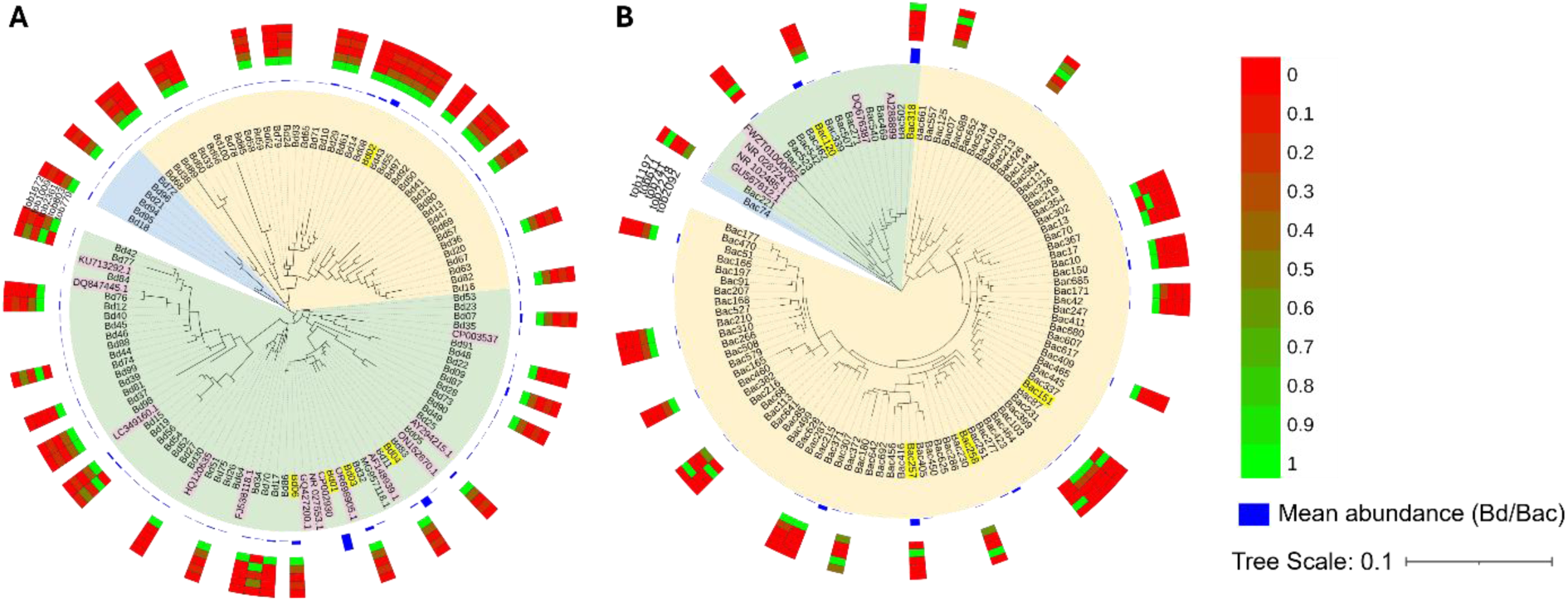
Maximum likelihood-based phylogenetic analysis of the 100 most abundant ASVs affiliated with the bacterial orders of the predators Bdellovibrionales (A) and Bacteriovoracales (B), based on 16S rRNA gene sequences. Distinct phylogenetic clades are color-coded. The five most abundant ASVs in each group are highlighted in yellow, and typed strains are indicated in pink. Outer rings represent the normalised (z-score) relative abundances of the top five gram-negative bacteria negatively correlated with predators (Separately for Bd and Bac) across soil samples, visualised as a heatmap (red = high, green = low). Blue bar denotes the mean abundance of each ASV of Bd and Bac. (Tree scale = 0.1).

### BALO dynamics

The dominant Bd and Bac ASVs collectively accounted for 41.2 ± 17.1% and 29.0 ± 3.85%, and 16.8 ± 2.54% and 26.0 ± 4.2% at the MF and OP sites, respectively. They were mostly shared between sites and treatments. Noteworthy, in MF field samples, in both manure and compost, Bd01 and Bd04 were abundant over the time series, with the latter increasing during the cooler seasons; In OP field samples, larger shifts in Bac ASV dominance were observed between time points and fertilization treatments (Figure 4, Table S11). Few significant potential associations between dominant Bd and Bac ASVs with GNtobs were detected using negative Kendall correlations: Bd02, Bd03, and Bd06 correlated with GNtob393 (Xanthomonadaceae), GNtob779 (Moraxellaceae), and GNtob803 (Yersiniaceae), respectively, and Bac318 with GNtob1197 (Gemmataceae) (Figure 5). The overlap between Bd- and Bac-associated GNtobs was consistently lower at the MF site than at the OP site under mineral, and more markedly, organic fertilization (Figure S10).

**Figure 4.**
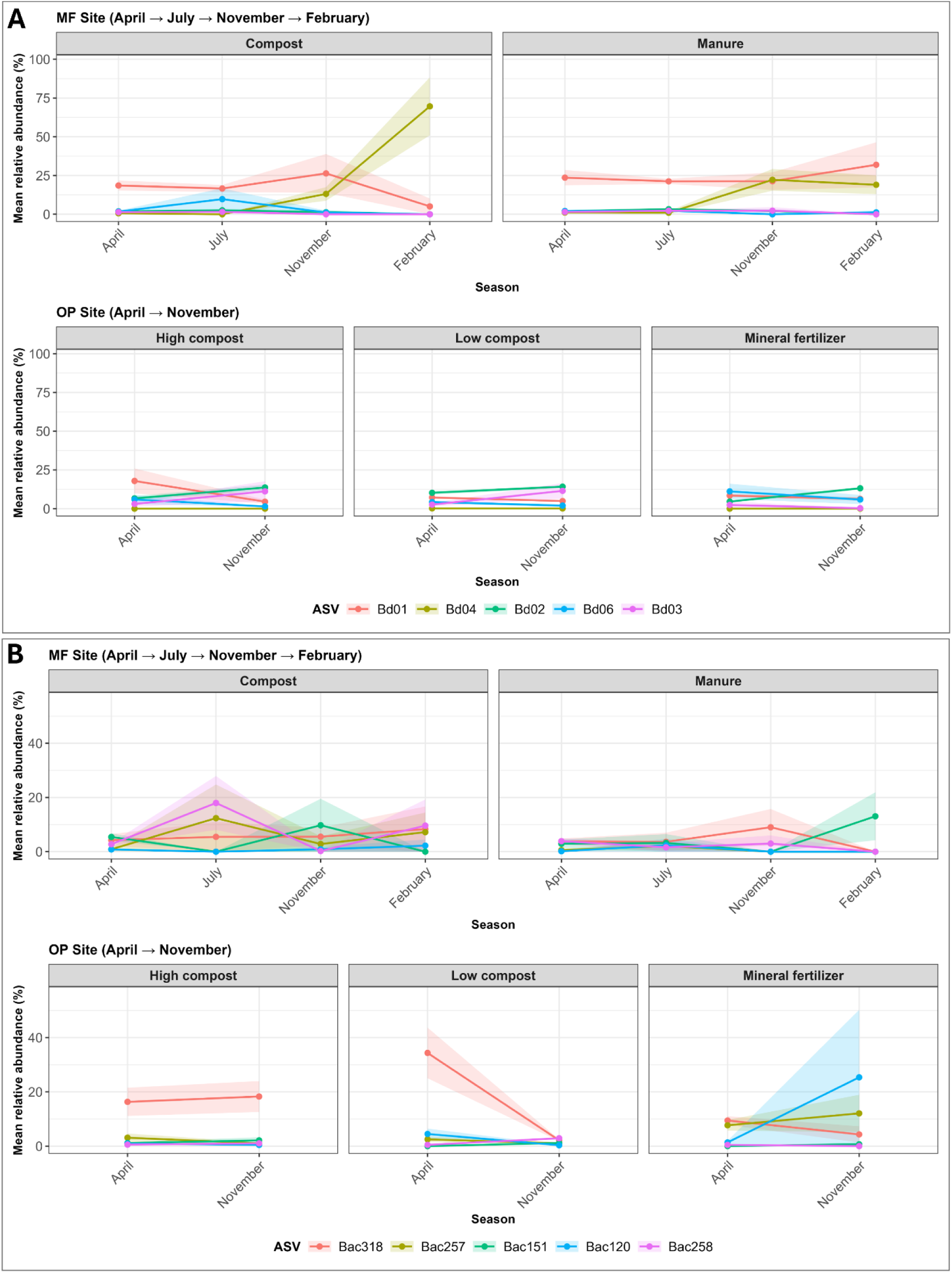
Seasonal dynamics of representative ASVs from Bd (Panel A) and Bac (Panel B) across fertilization treatments at the MF (top) and OP (bottom) sites. ASVs are the top 5 most abundant per treatment and the most consistently prevalent across seasonal samples.

**Figure 5.**
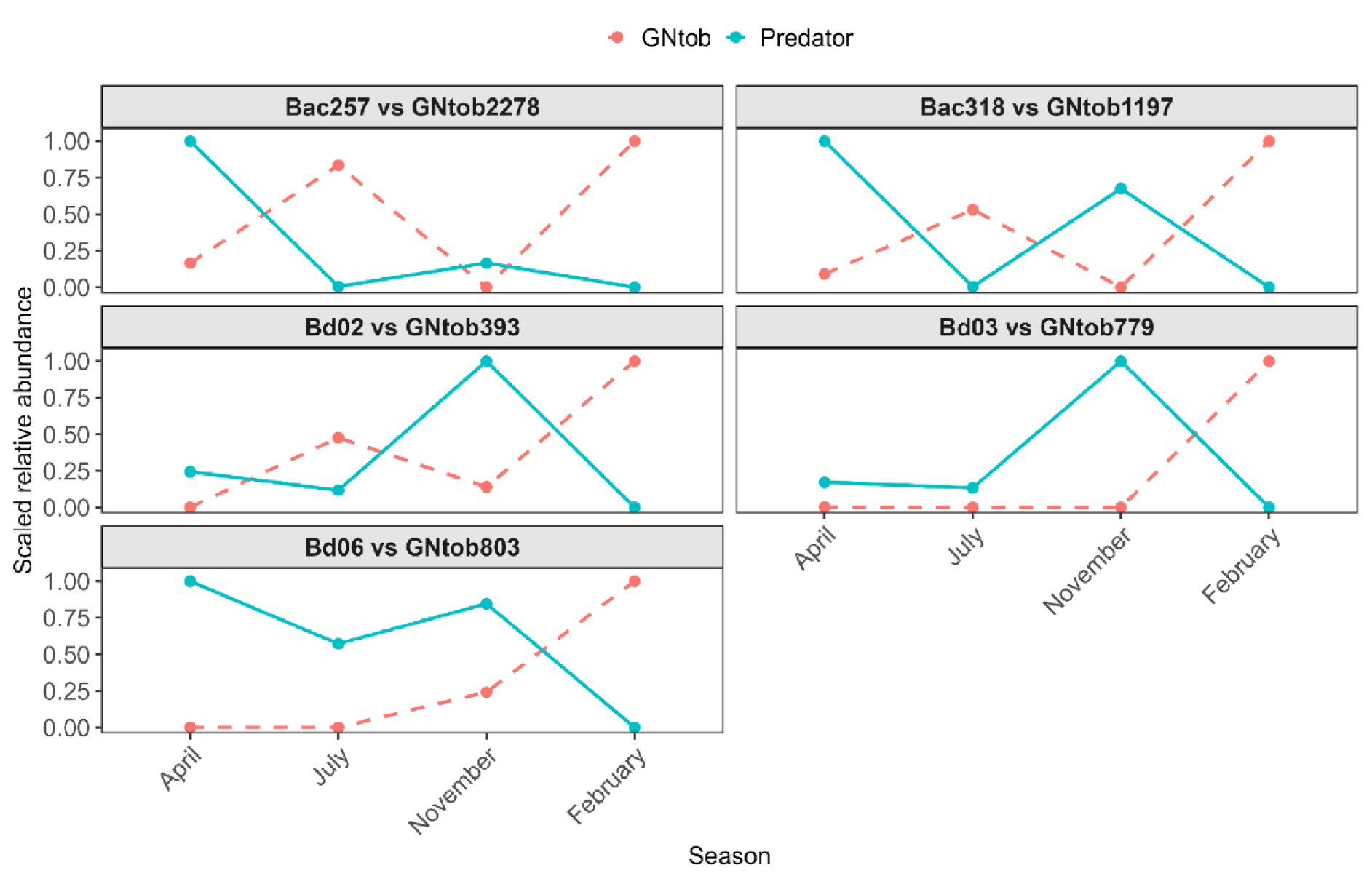
Seasonal dynamics of interactions between BALOs and gram-negative bacteria (GNtobs) in agricultural soil. (Kendall τ >-0.5, p< 0.05). *GNtob2278 = Beijerinckiaceae, GNtob1197 = Gemmataceae, GNtob779 = Moraxellaceae, GNtob393 = Xanthomonadaceae, GNtob803 = Yersiniaceae.

### Co-occurrence network analysis

At the MF site, co-occurrence networks in the compost and manure treatments exhibited low average degrees (2.57 and 2.65, respectively) (mean number of edges per node) and high modularity (0.74 - 0.78) (strength of division of a network into modules) suggesting sparsely connected but strongly compartmentalized network structures. Networks differed in graph density (0.021 vs 0.006) and diameter (6 vs 21), with the manure network showing a larger diameter and lower density, consistent with an expanded and weakly connected topology despite higher GNtob node richness (Figure 6A, B). At the OP site, networks of the fertilization treatments had higher connectivity, with average degree ranging from 8.7 to 10.46, along with tob node numbers (431-504) and larger predator node pools (up to 299 Bd and 234 Bac nodes). Modularity remained high but was consistently lower than at the MF site (0.49-0.64), while network diameter (11-14) and average path length (4.84-5.48) indicated more compact structures and interconnections compared to the MF field (Figure 6, Table S12).

**Figure 6.**
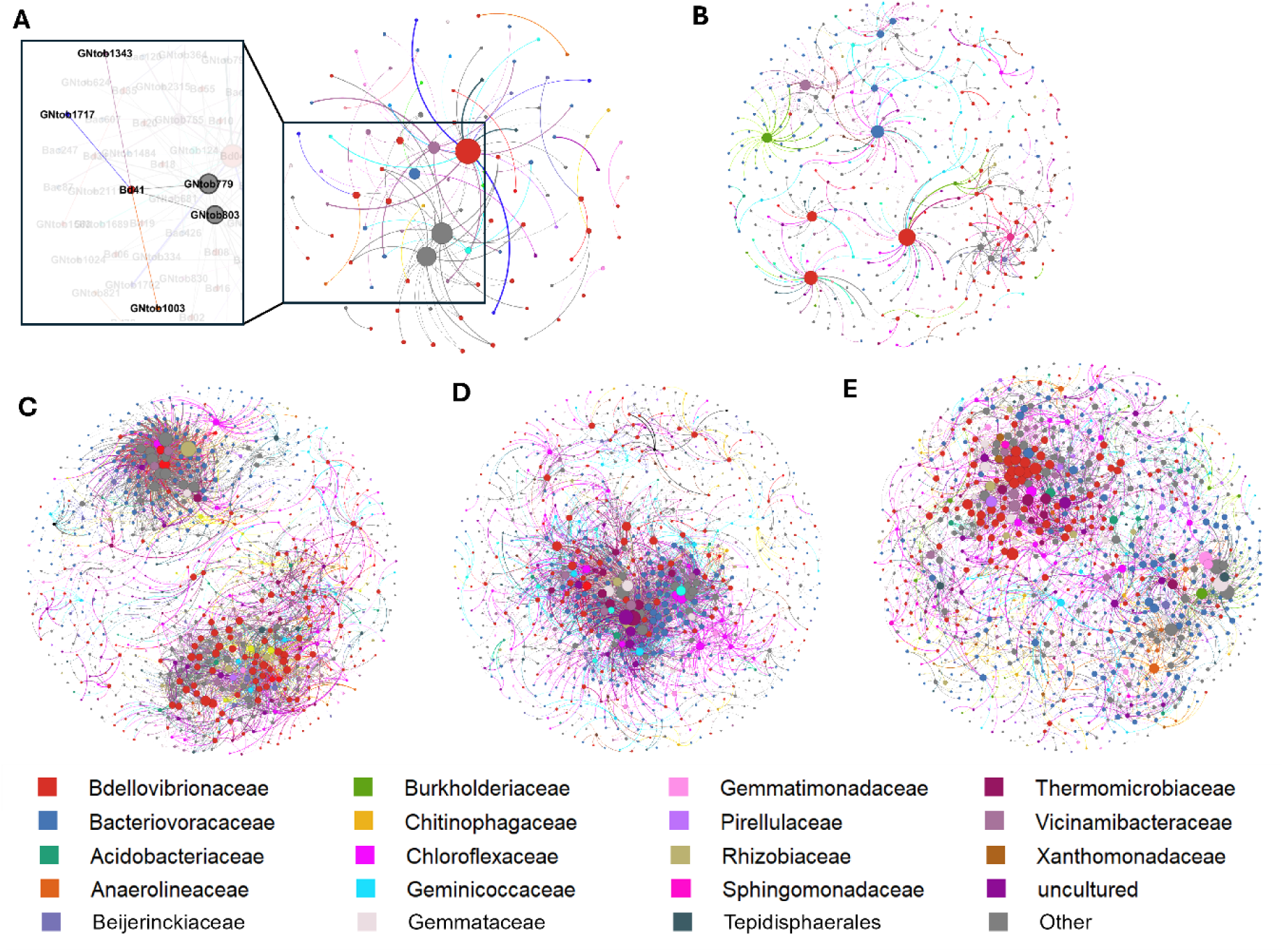
Negative co-occurrence networks (Kendall’s τ < -0.3, *p* < 0.05) between Bdellovibrionales (Bd) and Bacteriovoracales (Bac) predators and gram-negative tobs (GNtobs) across fertilization regimes at the Model Farm (MF) and Organic Plot (OP) sites. (A-B) Compost and manure treatments at the MF site; (C-E) mineral, low-compost, and high-compost treatments at the OP site. *Node colour denotes bacterial order (see legend), node size represents degree centrality, edge colour indicates target association, and edge thickness corresponds to correlation strength (|τ|). Inset in A: detail of the network extended to specific GNtobs.

## Discussion

This study uncovered a hitherto unknown diversity of BALOs that differentially respond to temporal and spatial factors, fertilization type and environmental parameters, and dynamically interact with gram-negative members the soil microbiome. These observations support niche differentiation between BALOs, which in turn may explain their observed diversity, and suggest cascading effects on soil bacterial community structure.

Based on QPCR, fertilization only marginally affected the sizes of the GEP, qBd and qBac communities, while season was a main driver of change. This, in spite of the different types, quantities, and time of application of the fertilizers. Contrary to all other samplings, GEP population sizes at the MF and OP sites were similar in April but different in November. This may be caused by ploughing the MF field after harvest, a practice known to reduce bacterial abundance ^52^. At all other sampling times, seasonal effects appeared to be decoupled from fertilization and from site in affecting GEP, qBd, and qBac community size. However, while sizes remained similar, fertilization treatment and site, along with seasons, explained part of the GEP β-diversities. In addition, we found that GEP composition was quite stable, with families (and genera) largely shared across treatments at each site. Although the fields were in close proximity (∼90–100 m), half of GEP ASVs differed between sites, probably due to taxon micro-diversity, suggesting distance-decay. Distance-decay was demonstrated to significantly influence community divergence even within single soil types and short distances ^52,53^. None of the GEP, Bd and Bac communities were directly influenced by carbon or nitrogen levels but in all, humidity, moisture and temperature were drivers of β-diversity. As in other studies of semi-arid regions, fertilization may act as a contributing but lesser factor of diversity than soil physical conditions ^54,55^. Along these lines, a study in rice paddies showed BALOs community structure to be very responsive to soil moisture and to potential BALO predator-prey interaction networks, but not to fertilization ^37^. Previous studies with *B. bacteriovorus* in soil microcosms also established that water connectivity of the soil pore network was central to sustain predator viability over time, and that its breakdown at low moisture levels enhanced prey refuges ^38,56^. Other studies established temperature to strongly impact BALO communities, directly and through shifts in prey ranges and predation ability ^33,38,57,58^. Taken together, physical parameters related to seasonal changes as well as to agricultural practice (irrigation) appear to act as major modifiers of the soil microbiome and of the BALO β-diversity in soil, reinforced in agricultural soil, by secondary factors like fertilization.

Numerous BALO ASVs clustered within phylogenetic clades that are barely represented by cultured isolates. 16S rRNA gene community sequencing and environmental metagenome analyses have revealed an extended BALO diversity in soil ^37,59,60^ and in aquatic ecosystems ^38,61^. Nonetheless, BALOs can be shared locally- here between the MF and OP sites- but also across large distances ^38^. An important limitation to interpreting these data in a biogeography context is that appraisal by ASVs or operational taxonomic units does not necessarily represent the genomic, physiological, and ecological diversity of the organisms, implying an even greater (micro)diversity. The BALO diversity observed here included both abundant ASVs present across treatments and rare, low-frequency lineages. Most dominant BALO ASVs substantially diverged from described cultured strains. For example, Bd02 and Bd06 together accounted for ∼24% of predator reads and diverged by >11% from the closest type strains (*Bdellovibrio bacteriovorus* HD100 and *B. exovorus* JSS), respectively. Noteworthy, these two species differ in predation strategy, the former penetrating its prey (periplasmic predation) and the latter consuming from the outside (epibiotic predation). These two modes of prey use may have ecological consequences, as their dynamics and predation efficiency may differ ^62^. Some ASVs, like Bd03, showed higher relative abundance under compost and manure treatments. Bd03 diverged by ∼10% from *B. bacteriovorus* Tiberius, a strain isolated from the Tiber River ^63^, a stream polluted by organic compounds. Bd01, an abundant ASV (∼21.6%), was loosely related to the soil isolate *B. bacteriovorus* HD100 (7% divergence) and was placed outside well-resolved reference clades ^64,65^. Similarly, within the Bacteriovoracales, dominant ASVs lacked close cultured representatives. Bac257 and Bac318, consistently detected under organic and manure fertilization diverged by 9 and 11% from *Halobacteriovorax marinus* SJ, and Bac151 diverged from *Peredibacter starrii* by ∼8% and did not cluster within established lineages. None of the dominant Bac ASVs were closely affiliated with *Bacteriovorax stolpii*, one of the few Bac terrestrial isolates. These findings align with previous reports of unexplored microbial divergence across soil ecosystems ^66,67^, and a large unknown BALO diversity ^38^.

Patterns of prevalence revealed two broad ecological strategies among BALO predators. Persistent strains, such as Bd02, Bac257, and Bac258, maintained relatively stable abundance across sites, treatments, and seasons, consistent with ecological tolerance and flexible prey use, including efficient exploitation of prey resources, while other strains appeared transiently. This contrast between persistent and opportunistic strategies may fit co-existence models in microbial predator-prey systems, where stability can arise from the balance between generalists and condition-dependent specialists ^68–70^. The interaction landscape was explored using Kendall negative correlations ^38^, i.e. associations exhibited inverse seasonal patterns, with GNtob abundance typically peaking when predator abundance was low. It would then be expected that persistent strains have large potential interaction ranges, and associate with numerous GNtobs. However, serious limitations arise when interpreting these relationships. A reason being the relatively long span between sampling times, more so at OP. Given the complexity of microbial population dynamics in soil, these patterns may only partly reflect predator-prey dynamics like kill-the-winner ^69,71,72^, and include other direct and indirect responses. Environmental genomics, stable-isotope tracing, and *in-situ* hybridization to track prey-predator interactions at the individual level would certainly help decipher and validate the contributions of BALOs to trophic networks in the environment ^32,73–76^ and more particularly *in-situ* prey range, under an experimental requisite of frequent sampling ^38^.

Nevertheless, co-occurrence network analysis revealed that fertilization and seasonal variations markedly influenced the interactions between BALOs and gram-negative members of the bacterial community. At the MF site, compost and manure networks displayed high modularity and sparse structure. At the OP site, predator diversity and connectivity remained comparatively stable across treatments, likely reflecting the cumulative effects of long-term management practices. Variables like soil management -including fertilization, crop cover, distance-decay, and numerous unaccounted for bottom-up and top-down factors ^77,78^ contribute to the heterogeneity of soil microhabitats (e.g. pore sizes, mineral and organic matter composition, suspended particles, biofilms, and else) ^68,72,77–82^ which in turn may facilitate predator and prey adaptations, including shifts in prey preference, growth in suspension or in biofilm phases, defense mechanisms, and more ^82,83^, and contribute to niche segregation between Bd and Bac, and between strains within these clades.

In conclusion, this study examined the effects of soil management of two agricultural fields on the soil microbiome over time, and more specifically on the diversity and dynamics of BALOs and their potential association networks. The observed response of BALO communities to moisture conditions confirms small scale microcosm experiments at the field level ^56,80^, and shifts due to fertilization regime may be a consequence of changes in prey abundance and composition ^83,84^. BALOs were shown to play a major role as top-down controllers of soil food webs under shifting resource levels ^32^. Whether these features are related to the large BALO diversity uncovered here, with varying prey ranges ^32^ thus enabling ecological plasticity of the predators’ community remains to be confirmed. These results of the diversity, dynamics and interactions of BALOs in agricultural fields contribute to the growing understanding that micro-predators provide important ecological functions that can greatly contribute to improve soil microbiome manipulations based on agronomic practices, including microbial inoculations to enhance beneficial, and sustainable environmental and agricultural outcomes ^27,85–87^.

## Acknowledgement

Part of this work was conducted at the Helmsley Model Farm for Sustainable Agriculture, located at Newe Ya’ar, the northern branch of the Volcani Center, Israel’s Agricultural Research Organization. The Model Farm was established in 2018 for studying, demonstrating and implementing sustainable agricultural practices. We thank Ronen Kfir for his on-site assistance during sampling at the Model Farm, and Itai Shulner for his help with sampling in the organic plot.

## Data availability

Demultiplexed sequencing data have been deposited in the Sequence Read Archive (https://www.ebi.ac.uk/) under accession numbers; PRJNA1265772, PRJNA1263579 and PRJNA1264548 for GEP, Bd and Bac, respectively.

## Consent for Publication

All authors have approved the final version of the manuscript and consent to its submission for publication.

## Ethics statement

No human beings or other animals were subjects of this study.

## Authors contributions

Alka Kumari performed sampling, analysis and parts of writing; Rolf Lood and Eddie Cytryn contributed to the methodology, to obtain funding and revised the manuscript; Ofra Matan provided technical support; Yael Laor and Gil Eshel were responsible for the field experiments at the Newe Ya’ar station and revised the manuscript

