## Supplementary Tables for "Site, Fertilization and Season Structure the Soil Microbiome and its Interactions with *Bdellovibrio* and Like Organisms Predators"

**Table S1.** Sample locations and soil parameters (Excel table S1).

**Table S2.** Primers used in this study.

| Primers | Sequences | Specificity | Use | PCR conditions | Ref |
|---|---|---|---|---|---|
| Bd_824F | ACTTGTTGTTGGA GGTAT | Bdellovibrionales (Bd) | Sequencing | 98°C/10s, 50°C/5s, 68°C/1s, 28 cycles | 1 |
| Bd_1222R | TTGTAGCACGTGT GTAG | | | | |
| Bx_341F | TACGGGAGGCAG CAG | Bacteriovoracales (Bac) | | 98°C/10s, 50°C/5s, 68°C/1s, 28 cycles | 1 |
| Bx_672R | TACCCCTACATGC GAAATTCC | | | | |
| 16S_515F | GTGCCAGCMGCC GCGG TAA | General prokaryote (GEP) | | 98°C/10s, 50°C/5s, 68°C/1s, 28 cycles | 2 |
| 16S_806R | GGACTACHVGGG TWTCTAAT | | | | |
| Bd_347F | GGAGGCAGCAGT AGGGAATA | Bdellovibrionaceae (qBd) | QPCR | 95°C/3min, 95°C/30s, 54°/35s, 40 cycles | 3 |
| Bd_549R | GCTAGGATCCCTC GTCTTACC | | | | |
| Bx_519F | CAGCAGCCGCGGT AATAC | Bacteriovoracaceae (qBac) | | 95°C/3min, 95°C/30s, 51°/35s, 40 cycles | 4 |
| Bx_677R | CGGATTTTACCCC TACATGC | | | | |
| 16S_1048F | GTGSTGCAYGGYT GTCGTCA | General prokaryote (GEP) | | 95°C/3min, 95°C/30s, 56°/30s, 40 cycles | 5 |
| 16S_1175R | ACGTCRTCCMCAC CTTCCTC | | | | |

**Table S3.** α-diversity estimates (Observed, Shannon, and Simpson) of General Prokaryotes at the MF and OP sites. Statistical comparison was performed using the Kruskal-Wallis test on the Shannon index across all treatment-time combinations ($\chi^2$ (13) = 29.26, p = 0.006).

| Site | Sampling time and treatment | Observed | Shannon | Simpson |
|---|---|---|---|---|
| MF | **April 2022, Compost** | 662.3 ± 16. | 5.8 ± 0.1 | 0.99 ± 0.00 |
| | **April 2022, Manure** | 664 ± 320.2 | 5.8 ± 1.3 | 0.99 ± 0.1 |
| | **July 2022, Compost** | 654 ± 41.0 | 6.0± 0.1 | 0.99 ± 0.00 |
| | **July 2022, Manure** | 664.8 ± 59.0 | 5.8 ± 0.1 | 0.99 ± 0.00 |
| | **November 2022, Compost** | 444 ± 4.0 | 5.5 ± 0.0 | 0.99 ± 0.00 |
| | **November 2022, Manure** | 373 ± 188.7 | 5.2 ± 0.6 | 0.98 ± 0.02 |
| | **February 2023, Compost** | 376.7 ± 24.5 | 3.8 ± 0.1 | 0.86 ± 0.00 |
| | **February 2023, Manure** | 620.4 ± 64.8 | 5.3 ± 0.2 | 0.96 ± 0.00 |
| OP | **April 2022, Mineral fertilizer** | 654.7 ± 20.2 | 5.8 ± 0.1 | 0.99 ± 0.00 |
| | **April 2022, Low compost** | 648 ± 14.9 | 5.7 ± 0.0 | 0.99 ± 0.00 |
| | **April 2022, High compost** | 658.7 ± 26.0 | 5.8 ± 0.1 | 0.99 ± 0.00 |
| | **November 2022, Mineral fertilizer** | 502 ± 27.3 | 5.5 ± 0.1 | 0.99± 0.00 |
| | **November 2022, Low compost** | 458.7 ± 10.1 | 5.5 ± 0.0 | 0.99± 0.00 |
| | **November 2022, High compost** | 478.7 ± 13.8 | 5.5 ± 0.0 | 0.99 ± 0.00 |

Between sites (Shannon index: W = 383, p = 0.993; Simpson index: W = 380, p = 0.954; Observed richness: W = 429, p = 0.461, Wilcoxon rank-sum test; Between fertilization treatments (Shannon index: $\chi^2(4) = 0.27$, p = 0.992; Simpson index: $\chi^2(4) = 0.79$, p = 0.94; Observed: $\chi^2(4)=0.7$, p = 0.951; Kruskal-Wallis test); Between seasons (Shannon index: $\chi^2(3) = 27.22$, p < 5.28e-06; Simpson index: $\chi^2(3) = 21.33$, p = 9e-05; Observed: $\chi^2(3) =27.94$, p < 3.73e-06; Kruskal-Wallis test).

**Table S4.** α-diversity estimates (Observed, Shannon and Simpson) of Bdellovibrionales at the MF and OP sites. Statistical comparison was performed using the Kruskal-Wallis test on the Shannon index across all treatment-time combinations ($\chi^2$ (13) = 25.24, p = 0.021).

| Site | Sampling time and treatment | Observed | Shannon | Simpson |
|---|---|---|---|---|
| MF | **April 2022, Compost** | 91 ± 3.8 | 3.8 ± 0.1 | 0.95 ± 0.01 |
| | **April 2022, Manure** | 92.2 ± 4.7 | 3.6 ± 0.2 | 0.92 ± 0.03 |
| | **July 2022, Compost** | 76.3 ± 5.7 | 3.7 ± 0.1 | 0.93 ± 0.01 |
| | **July 2022, Manure** | 90.4 ± 3.4 | 3.8 ± 0.1 | 0.94 ± 0.01 |
| | **November 2022, Compost** | 53 ± 21.7 | 3.0 ± 0.7 | 0.86 ± 0.08 |
| | **November 2022, Manure** | 33.2 ± 9.4 | 2.4 ± 0.2 | 0.82 ± 0.03 |
| | **February 2023, Compost** | 10.3 ± 7.9 | 1.1 ± 0.7 | 0.42 ±0.24 |
| | **February 2023, Manure** | 32 ± 14.2 | 2.2 ± 0.6 | 0.73 ± 0.13 |
| OP | **April 2022, Mineral fertilizer** | 73.5 ± 5.6 | 3.5 ± 0.2 | 0.95 ± 0.01 |
| | **April 2022, Low compost** | 87.7 ± 4.7 | 3.8 ± 0.1 | 0.96 ± 0.00 |
| | **April 2022, High compost** | 79.7 ± 9.0 | 3.5± 0.3 | 0.92 ± 0.04 |
| | **November 2022, Mineral fertilizer** | 132.5 ± 6.9 | 3.7 ± 0.1 | 0.94 ± 0.01 |
| | **November 2022, Low compost** | 123.7 ± 11.2 | 3.6± 0.2 | 0.93 ± 0.01 |
| | **November 2022, High compost** | 127.7 ± 5.7 | 3.6 ± 0.1 | 0.94 ± 0.02 |

Between sites (Shannon index: W = 275, p = 0.072; Simpson index: W = 218, p = 0.0055; Observed richness: W = 145.5, p = $8.1 \times 10^{-5}$, Wilcoxon rank-sum test); Between fertilization treatments (Shannon index: $\chi^2(4) = 3.5$, p = 0.478; Simpson index: $\chi^2(4) = 8.1$, p = 0.088; Observed: $\chi^2(4) = 15.96$, p = 0.003; Kruskal–Wallis test; Between seasons (Shannon index: $\chi^2(3) = 15.45$, p = 0.0015; Simpson index: $\chi^2(3) = 14.08$, p = 0.0028; Observed: $\chi^2(3) = 16.36$, p = 0.00096; Kruskal–Wallis test).

**Table S5.** α-diversity estimates (Observed, Shannon and Simpson) of Bacteriovoracales at the MF and OP sites. Statistical comparison was performed using the Kruskal-Wallis test on the Shannon index across all treatment-time combinations ($\chi^2$ (13) = 35.33, p = 0.001).

| Site | Sampling time and treatment | Observed | Shannon | Simpson |
|---|---|---|---|---|
| MF | April 2022, Compost | 91.7 ± 28.7 | 4.04 ± 0.2 | 0.98 ± 0.00 |
| | April 2022, Manure | 101.8 ± 30.4 | 3.9 ± 0.3 | 0.96 ± 0.01 |
| | July 2022, Compost | 6.7 ± 2.3 | 1.7 ± 0.3 | 0.79 ± 0.05 |
| | July 2022, Manure | 20 ± 6.0 | 2.7 ± 0.3 | 0.91 ± 0.02 |
| | November 2022, Compost | 31.3 ± 21.6 | 2.5 ± 0.7 | 0.85 ± 0.09 |
| | November 2022, Manure | 23.8 ±15.2 | 2.36 ± 0.5 | 0.86 ± 0.05 |
| | February 2023, Compost | 9.7± 5.8 | 1.7 ± 0.7 | 0.71 ± 0.17 |
| | February 2023, Manure | 35.8 ± 13.9 | 2.9 ± 0.5 | 0.89 ± 0.07 |
| OP | April 2022, Mineral fertilizer | 195.7 ± 16.0 | 4.1 ± 0.1 | 0.96 ± 0.01 |
| | April 2022, Low compost | 103.5 ± 36.1 | 3.0 ± 0.7 | 0.83 ± 0.08 |
| | April 2022, High compost | 198.5 ± 23.2 | 4.1 ± 0.2 | 0.95 ± 0.02 |
| | November 2022, Mineral fertilizer | 13.5 ± 6.7 | 1.9 ± 0.7 | 0.66 ± 0.22 |
| | November 2022, Low compost | 186.5 ± 6.2 | 4.6 ± 0.0 | 0.98 ± 0.00 |
| | November 2022, High compost | 171.2 ± 19.3 | 3.8 ± 0.2 | 0.93± 0.02 |

Between sites (Shannon index: W = 211, p = 0.0037; Simpson index: W = 313, p = 0.245; Observed richness: W = 132, p = $3.1 \times 10^{-5}$, Wilcoxon rank-sum test; Between fertilization treatments (Shannon index: $\chi^2(4)$ = 11.94, p = 0.018; Simpson index: $\chi^2(4)$ = 3.41, p = 0.491; Observed: $\chi^2(4)$ = 21.51, p = 0.001; Kruskal–Wallis test); Between seasons (Shannon index: $\chi^2(3)$ = 13.75, p = 0.00326; Simpson index: $\chi^2(3)$ = 6.77, p = 0.0798; Observed: $\chi^2(3)$ = 17.46, p = 0.00057; Kruskal–Wallis test).

**Table S6.** Permutation-based ANOVA of RDA between GEP community composition and environmental variables.

| Variable | Df | Variance | F value | Pr(>F) |
|---|---|---|---|---|
| Air temperature | 1 | 0.019 | 3.51 | 0.001 |
| Air humidity | 1 | 0.013 | 2.4 | 0.003 |
| TN | 1 | 0.008 | 1.55 | 0.067 |
| TC | 1 | 0.008 | 1.56 | 0.072 |
| pH | 1 | 0.007 | 1.3 | 0.161 |
| Soil moisture | 1 | 0.018 | 3.32 | 0.001 |
| Residual | 49 | 0.268 | NA | NA |

**Table S7.** Permutation-based ANOVA of RDA between Bdellovibrionales community composition and environmental variables.

| Variable | Df | Variance | F value | Pr(>F) |
|---|---|---|---|---|
| **Air temperature** | 1 | 0.037 | 4.98 | 0.001 |
| **Air humidity** | 1 | 0.016 | 2.19 | 0.021 |
| **TN** | 1 | 0.008 | 1.19 | 0.242 |
| **TC** | 1 | 0.009 | 1.32 | 0.179 |
| **pH** | 1 | 0.014 | 1.84 | 0.058 |
| **Soil moisture** | 1 | 0.017 | 2.29 | 0.015 |
| **Residual** | 49 | 0.368 | NA | NA |

**Table S8.** Permutation-based ANOVA of RDA between Bacteriovoracales community composition and environmental variables.

| Variable | Df | Variance | F value | Pr(>F) |
|---|---|---|---|---|
| **Air temperature** | 1 | 0.015 | 1.26 | 0.13 |
| **Air humidity** | 1 | 0.017 | 1.41 | 0.048 |
| **TN** | 1 | 0.015 | 1.19 | 0.296 |
| **TC** | 1 | 0.014 | 1.18 | 0.308 |
| **pH** | 1 | 0.019 | 1.63 | 0.013 |
| **Soil moisture** | 1 | 0.022 | 1.85 | 0.003 |
| **Residual** | 49 | 0.594 | NA | NA |

**Table S9**. Core microbiome community (GEP, Bd and Bac) across treatments and seasons (Excel table S9).

**Table S10**. Core microbiome community (GEP, Bd and Bac) between sites (Excel table S10).

**Table S11.** Relative abundance (%) of top 5 abundant BALOs (Bd and Bac) ASVs (Excel table S11).

**Table S12.** BALOs - GNtobs network correlation metrics at the MF and OP sites (Excel table S12).
