## Supplementary Figure for "Site, Fertilization and Season Structure the Soil Microbiome and its Interactions with *Bdellovibrio* and Like Organisms Predators"

**Supplementary Figures**

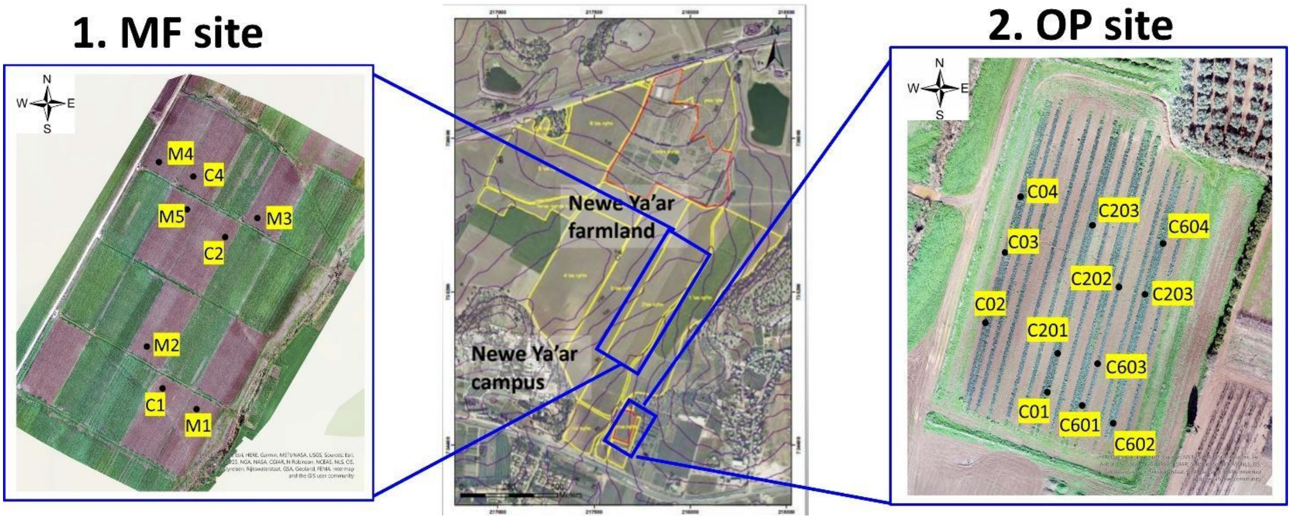


**Figure S1.** Spatial layout of experimental plots at the Newe Ya'ar Research Center. The Model Farm (MF) site83 layout (left), includes compost (C1, C2, C3, C4) and manure (M1, M2, M3, M4) treatment plots. The Organic Plot (OP) site84 layout (right) includes mineral fertilization (C0); low compost dose -10 m3.ha-1 (C20) and high compost dose - 60 m3.ha-1 (C60). Samples at the OP site were taken from plots where weed control was based on cultivation only.

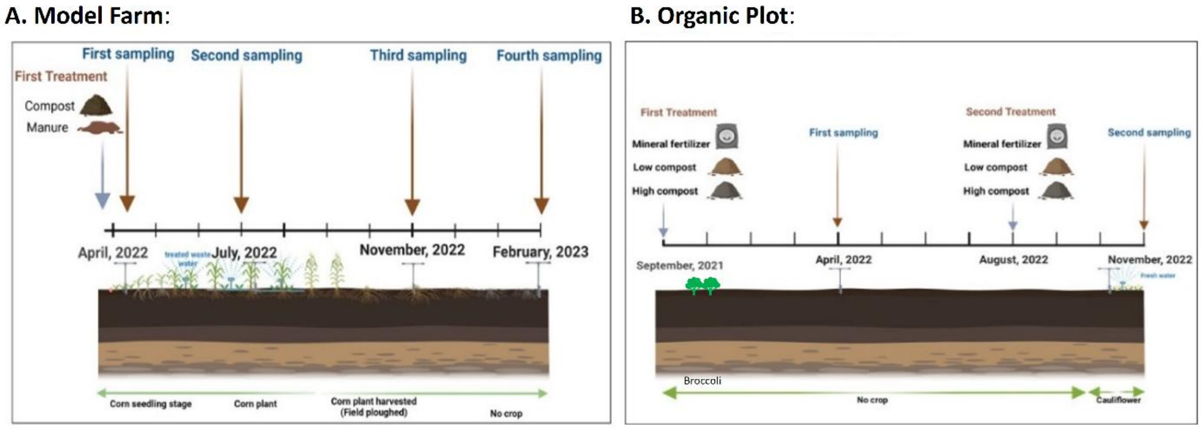


**Figure S2.** Treatments and sampling times. Soil samples were retrieved at the indicated dates from (A) Model Farm (MF) plots treated with (i) manure-based compost (35 m3.ha-1) derived from livestock manure and slaughterhouse waste residues (including blood, feces, and residual animal matter) and (ii) pre-stabilized cow manure (22 m3 .ha-1) from a dairy farm. (B) Organic Plot (OP) treated with mineral fertilizers (NPK at 200, 50, and 100 kg $ha^{-1}$) or with compost at high (60 $m^3$ $ha^{-1}$) or low (10 $m^3$ $ha^{-1}$) application rates after summer 2021 and in August 2022. Sampling was performed at four time points at the MF site (April, July, and November 2022, and February 2023) and at two time points at the OP site (April and November 2022).

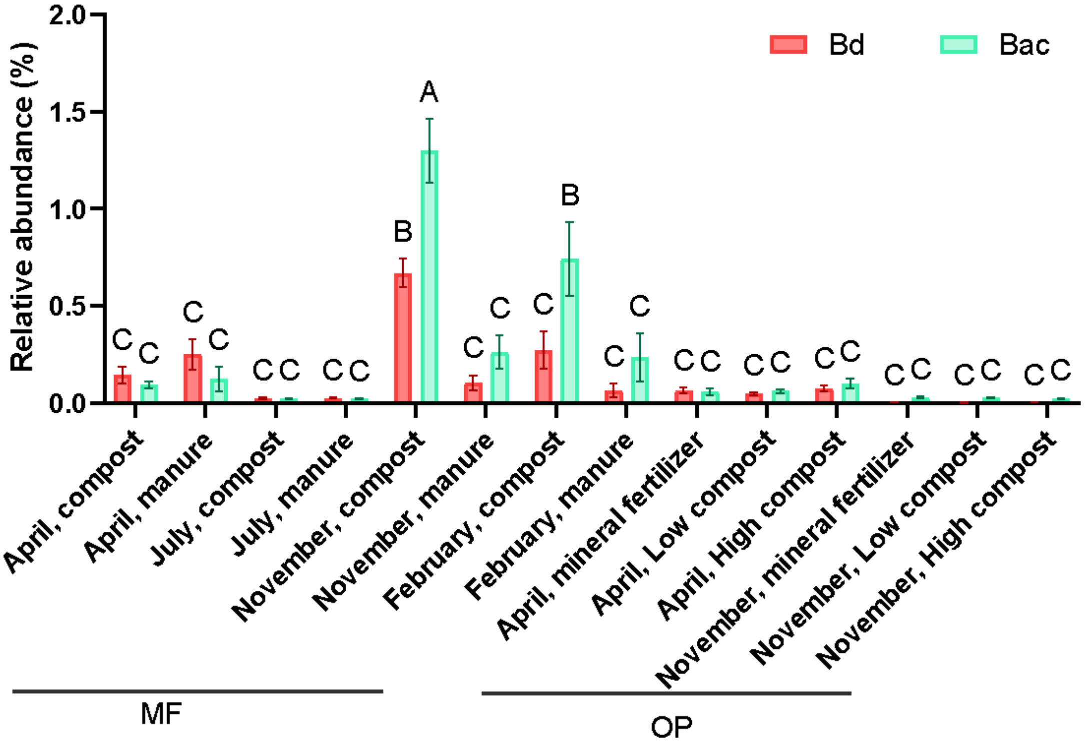


**Figure S3.** (A) Relative abundances Bdellovibrionaceae (Bd) and Bacteriovoracaceae (Bac). Error bars represent standard error. Different letters indicate statistically significant differences ($p < 0.05$, ANOVA with post hoc Tukey's HSD test): Lowercase Latin letters (a, b, c) for Bd, upper letters (A, B, C) for Bac. Statistical tests were performed separately for each group.

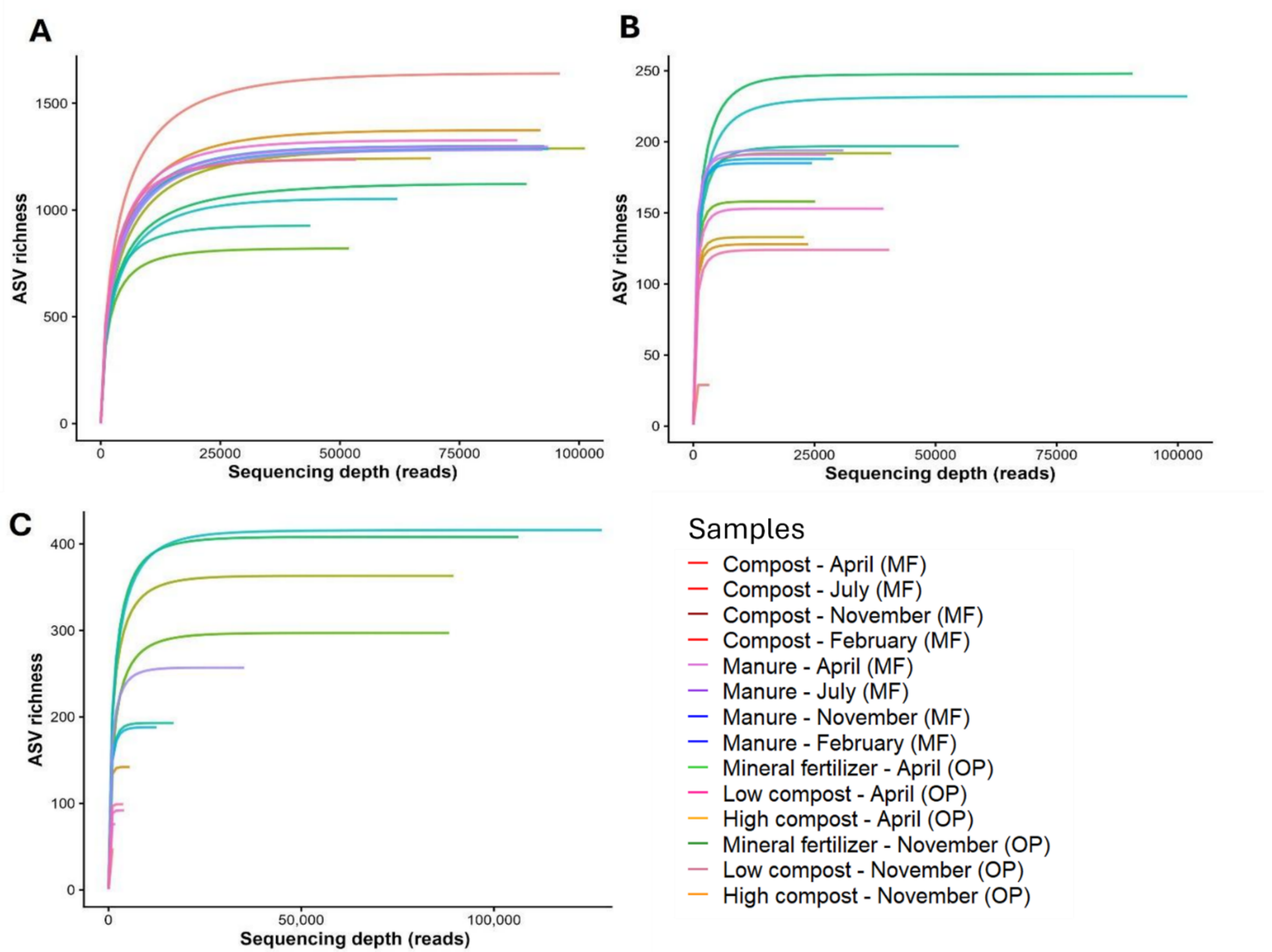


**Figure S4**. Rarefaction curves of ASVs vs processed 16S rRNA gene reads for General Prokaryote (A), (A), Bdellovibrionales (Bd) (B), and Bacteriovoracales (Bac) (C).

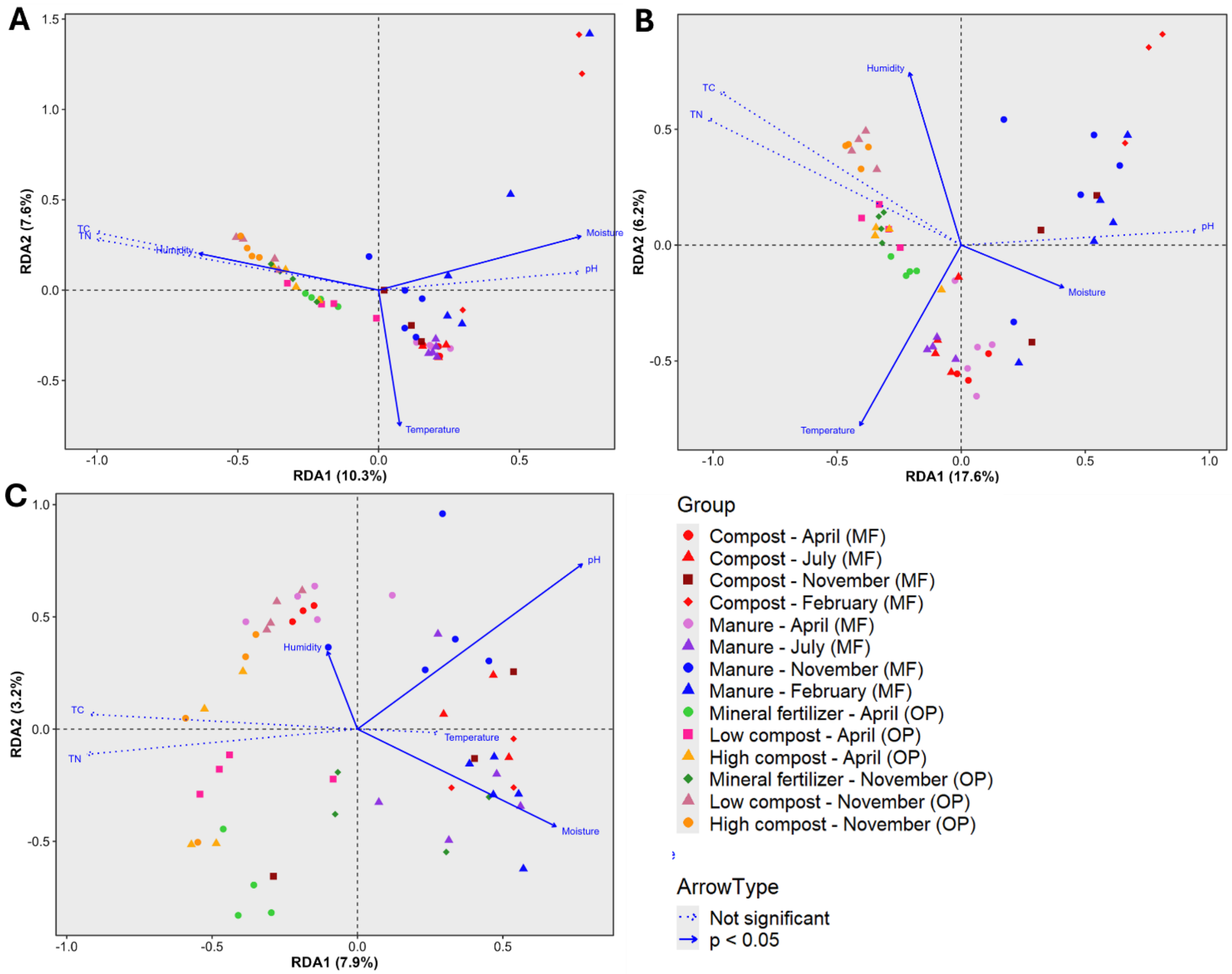


**Figure S5**. Redundancy Analysis plot of General prokaryote (A), Bdellovibrionales (B), and Bacteriovoracales (C) community composition in relation to environmental factors. Colours code for fertilization treatment and shapes, seasons. Solid arrows denote significant relationships ($p < 0.05$) and dotted arrows indicate non-significant variables.

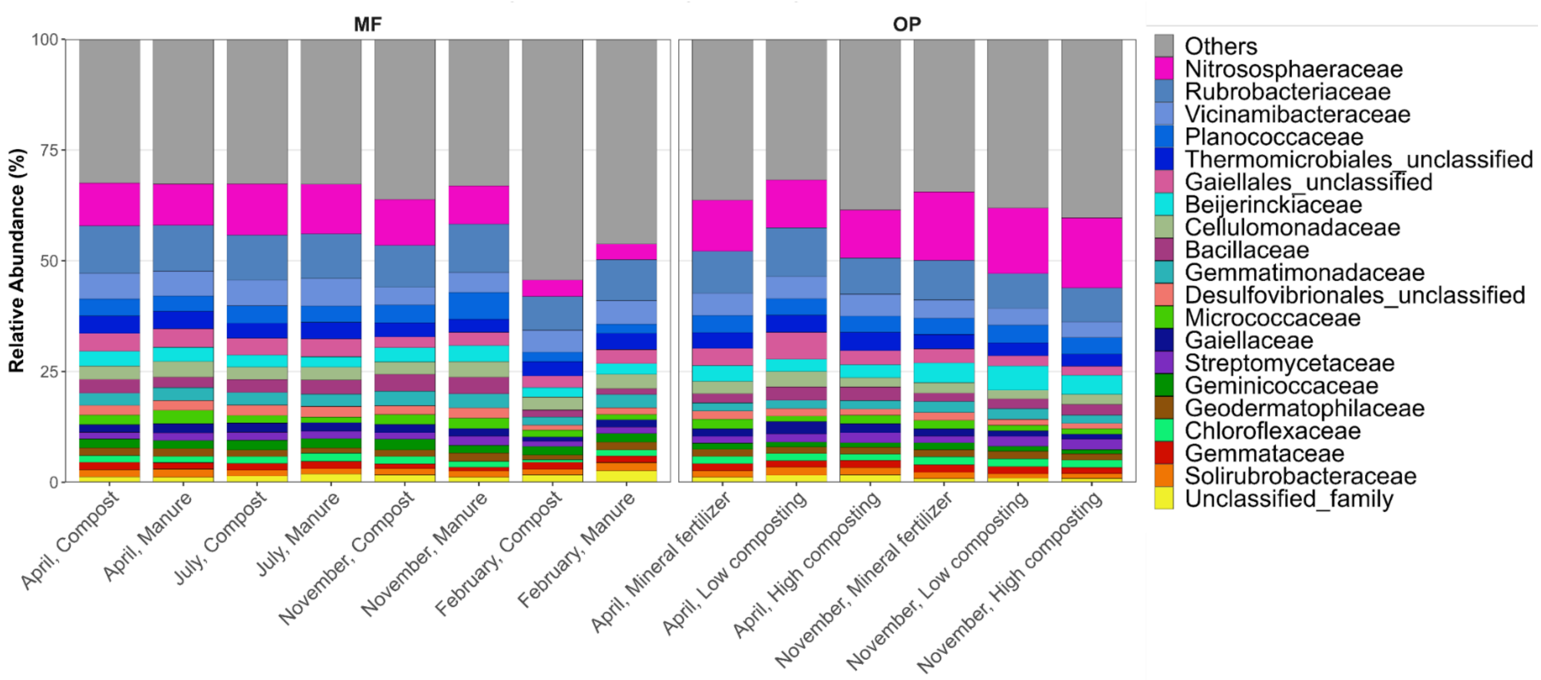


**Figure S6.** Relative abundance (%) of the General Prokaryotes community at the family level. Each bar represents the samples grouped by site (MF; OP) fertilization treatment, and sampling time.

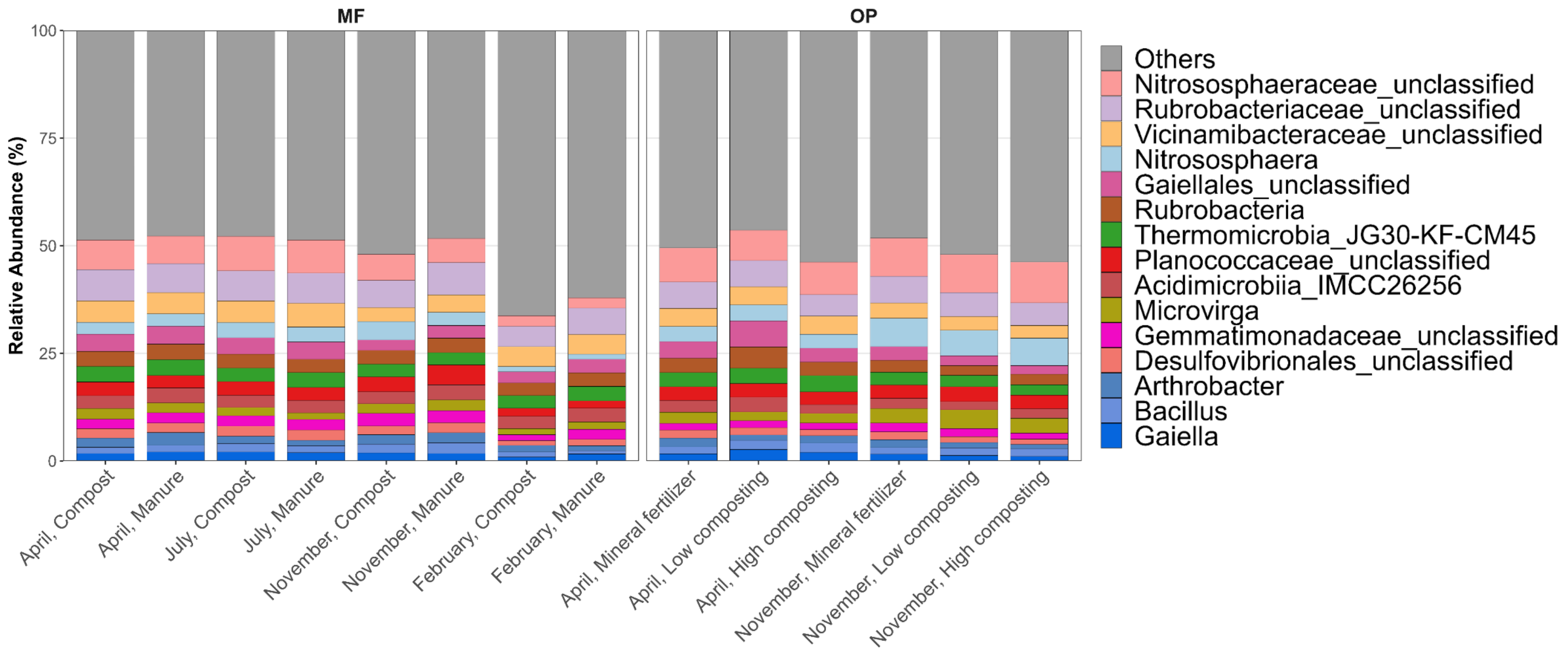


**Figure S7.** Relative abundance (%) of the General Prokaryote community at the genus level. Each bar represents the samples grouped by site (MF; OP) fertilization treatment, and sampling time.

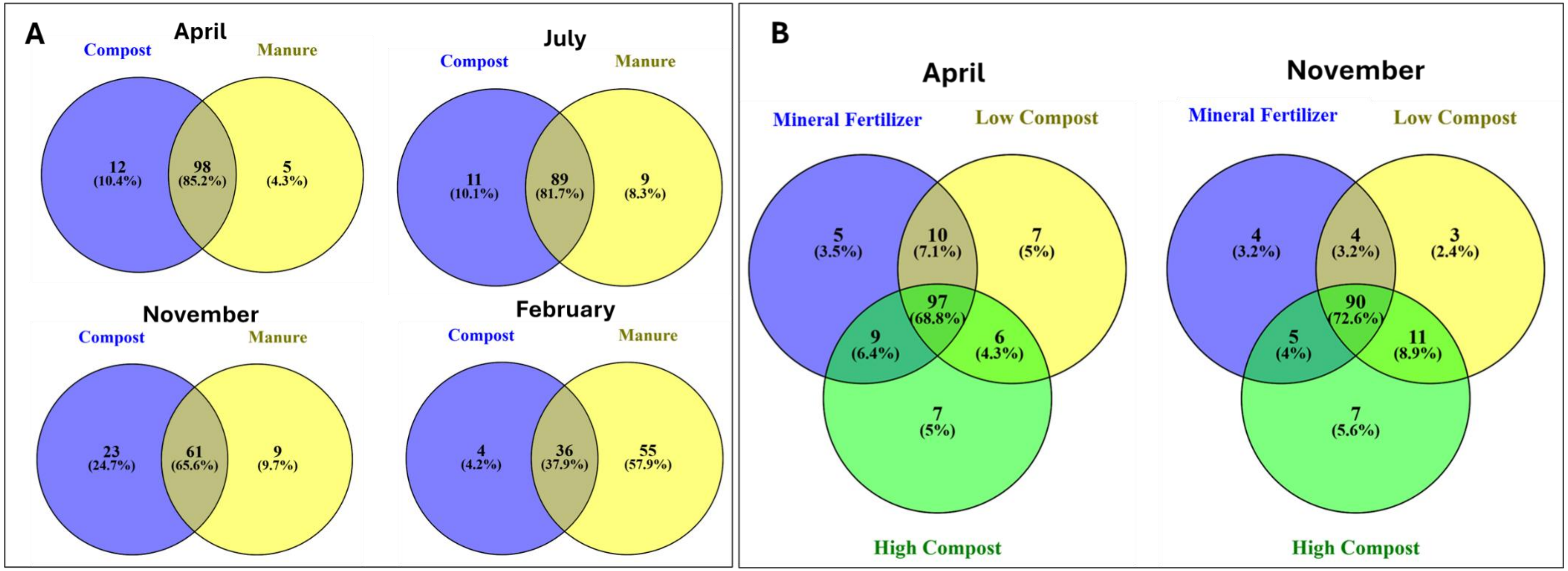


**Figure S8.** Venn diagrams of unique and shared fractions of core GEP families at the MF site (A), and at the OP site (B).

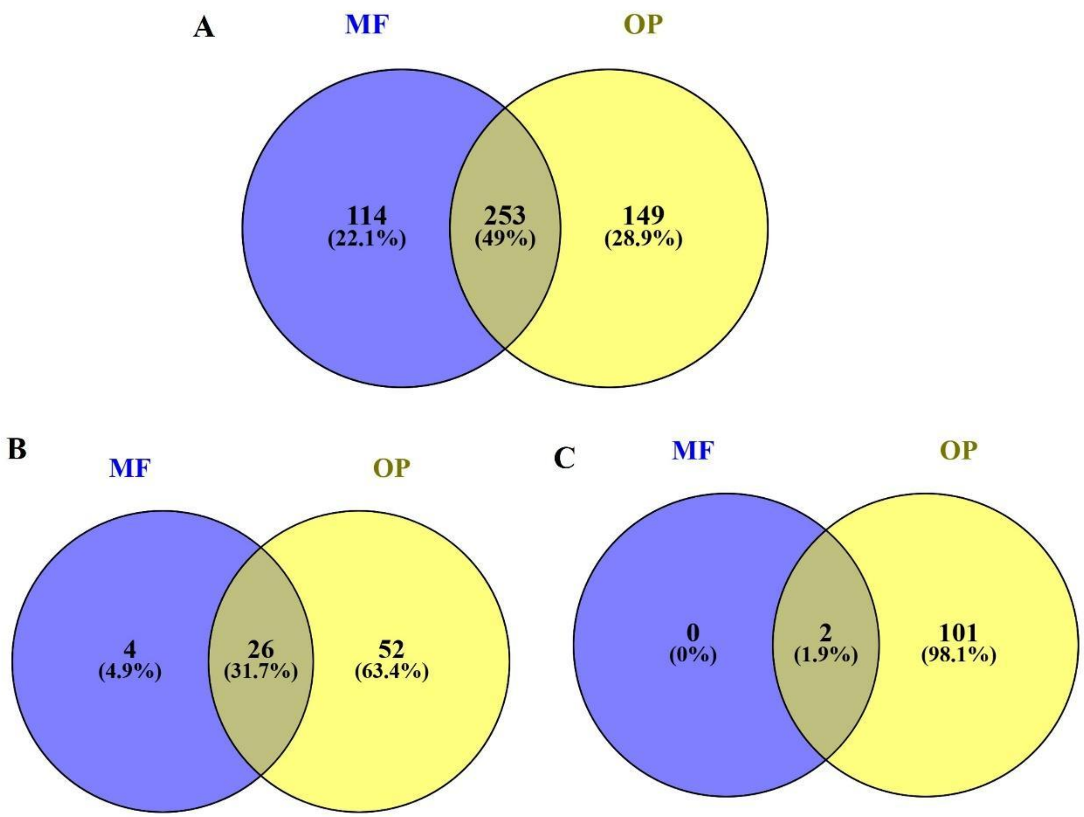


**Figure S9.** Venn diagrams of unique and shared fractions of core GEP, Bd and Bac ASVs at the MF site (A), and at the OP site (B).

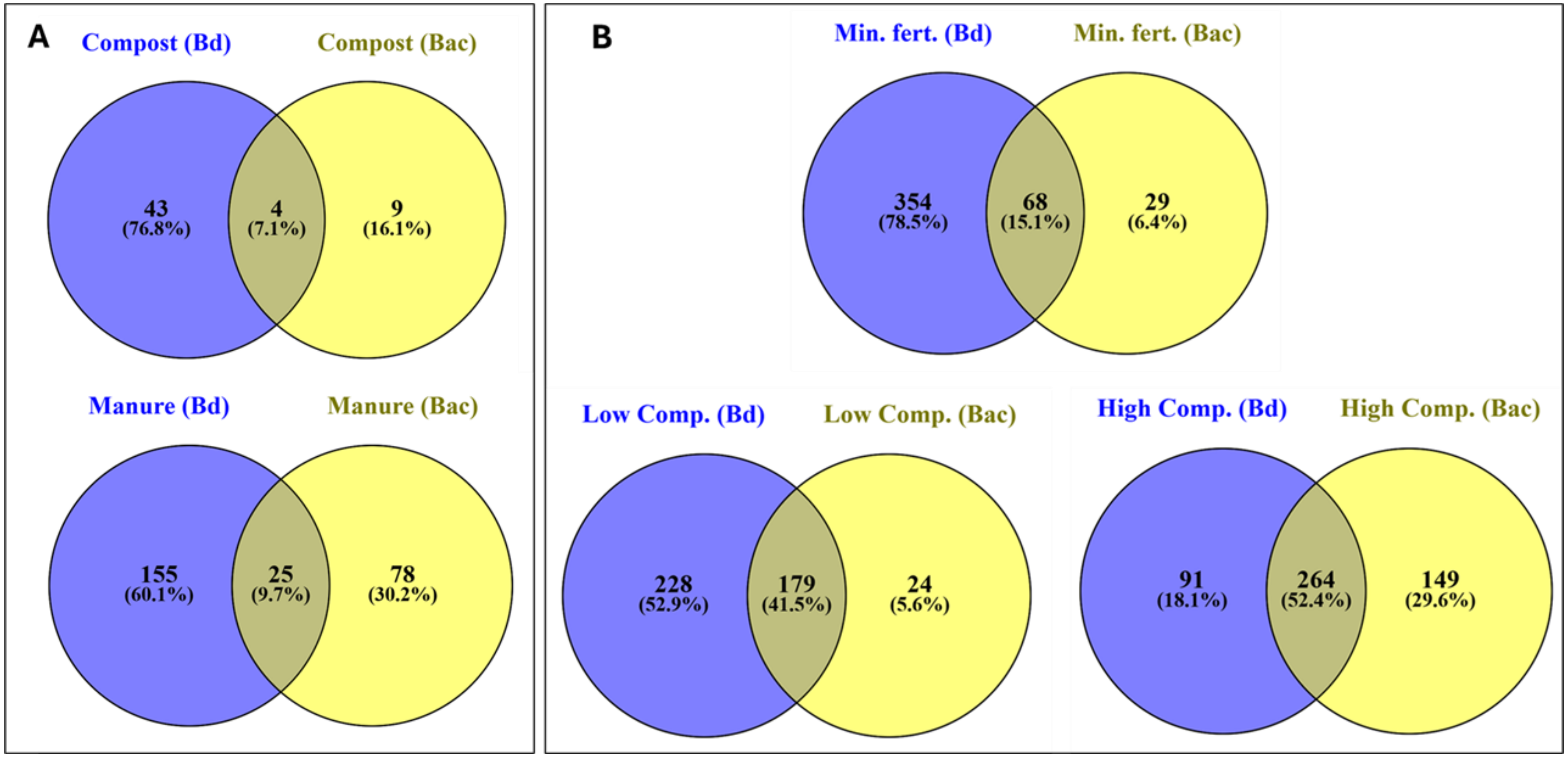


**Figure S10.** Venn diagrams of the unique and shared fractions of ASVs of Gram-negative bacteria (GNtobs) interacting with Bd and Bac ASVs in the different fertilization treatments, at the MF (A) and the OP (B) sites.
